# A synthetic ancestral Kinesin-13 depolymerises microtubules faster than any natural depolymerising kinesin

**DOI:** 10.1101/2022.04.11.487848

**Authors:** Hannah R. Belsham, Hanan M. Alghamdi, Nikita Dave, Alexandra J. Rathbone, Bill Wickstead, Claire T. Friel

## Abstract

The activity of a kinesin is largely determined by the ~350 residue motor domain and this region alone is sufficient to classify a kinesin as a member of a particular family. The Kinesin-13 family are a group of microtubule depolymerising kinesins and are vital regulators of microtubule length. Members of this family are critical to spindle assembly and chromosome segregation in both mitotic and meiotic cell division and play crucial roles in cilium length control and neuronal development. To better understand the evolution of microtubule depolymerisation activity in the Kinesin-13 family, we created a synthetic ancestral Kinesin-13 motor domain. This phylogenetically-inferred ancestral motor domain is the sequence predicted to have existed in the common ancestor of the Kinesin-13 family. Here we show that the ancestral Kinesin-13 motor depolymerises stabilised microtubules faster than any previously tested depolymerase. This potent activity is more than an order of magnitude faster than the most highly studied Kinesin-13, MCAK and allows the ancestral Kinesin-13 to depolymerise doubly-stabilised microtubules that are unaffected by MCAK. These data suggest that the ancestor of the Kinesin-13 family was a ‘super depolymeriser’ and that members of the Kinesin-13 family have evolved away from this extreme depolymerising activity to provide more controlled microtubule depolymerisation activity in extant cells.

## Introduction

The kinesin superfamily is a group of molecular motors that interact with the microtubule cytoskeleton. Kinesins are found in all eukaryotes, with 45 kinesin genes present in the human genome^1^. The characteristic ~350 residue motor domain marks a protein as a member of the kinesin superfamily. The kinesin motor domain is capable of a range of activities with respect to microtubules which, combined with the diversity of non-motor regions found within the superfamily, allow kinesins to carry out a wide array of cellular functions. Kinesin-Microtubule systems underpin many vital cellular processes such as cell division, cargo transport, development and maintenance of cell polarity, growth of neuronal processes and growth and function of cilia^2–8^.

Phylogenetic analysis of the kinesin superfamily has identified 17 individual families, many of which can be further divided into subfamilies^9, 10^. The activity of each kinesin family is largely determined by the motor domain, and this region alone is sufficient to classify most kinesin proteins to a family. However, how the structurally highly conserved motor domain can accommodate different activities – ranging from highly processive directed movement on the microtubule^11^ to non-directional short-lived diffusive motion^12^, and from promoting microtubule depolymerisation^13, 14^ to enhancing microtubule growth^15^ – is not well understood. Mutational studies on individual kinesins from different families provide some insight into the residues and regions of structure in the motor domain that tailor a specific kinesin to particular activities^16–19^. Whilst such studies have provided valuable knowledge, the extent to which they are generalisable to other motors in a subgroup or representative of the behaviour of a whole family is often unclear. This issue is exacerbated as data are often generated from a limited set of species, making the general applicability of findings to the activity of other motor domains, or to motors in phylogenetically less related organisms, difficult to ascertain. As a comprehensive approach to understanding how the motor domain sequence specifies the molecular activity of a kinesin family, we designed two reference sequences – the ‘consensus’ and the ‘ancestral’ sequences – for the motor domain of a kinesin family (Kinesin-13) using a set of phylogenetically diverse eukaryotes. The ‘consensus’ motor is a sequence that exists nowhere in nature but represents a hypothetical average molecule containing the most common residues at each position across the kinesin family. The ‘ancestral’ motor is the sequence inferred as most likely to have existed in the last common ancestor of all organisms that now possess sequences belonging to the specific kinesin family.

The Kinesin-13 family are a set of specialist microtubule depolymerases, with a centrally located motor domain flanked by divergent N- and C- terminal regions^20^. Kinesin-13s are major regulators of microtubule length and are particularly important during cell division^21–23^. The family is divided into 3 ancient subfamilies (denoted A, B and C). Subfamily B contains the most highly studied members of the family, including the mammalian Kinesin-13s KIF2A, KIF2B, and MCAK/KIF2C, although it is restricted to animals and their close relatives^10^. In contrast, Kinesin-13A (including human KIF24) dates back to the last common eukaryotic ancestor and is the most widely distributed family. Kinesin-13s are thought to drive depolymerisation by stabilising a curved conformation of the microtubule subunit, the α/β-tubulin heterodimer, resulting in destabilisation of the microtubule. Structures of the motor domain of either KIF2A or MCAK in complex with the α/β-tubulin heterodimer, show that they promote curvature of tubulin larger than that observed for similar complexes of tubulin with a Kinesin-1 motor domain^24–26^. The most highly studied Kinesin-13, MCAK, has no translocating activity, but diffuses along the microtubule without directional bias^12^. The ability of MCAK to stabilise curved conformation of tubulin is targeted to the microtubule end due to an atypical ATPase cycle in which only microtubule ends maximally accelerate ATP turnover^27^.

By creation of reference motor domain sequences, we sought to capture the activity associated with the Kinesin-13 family as a whole. The conceptual differences between the reference motors are non-trivial: the ancestral motor must have been functional but was not necessarily tuned to the specialised roles seen in extant cells. In contrast, the consensus motor could be either an optimal sequence for a family or a non-functional average. We expressed both the consensus and ancestral Kinesin-13 reference motor domains and characterised their behaviour with respect to microtubules. Here we show that, whilst the consensus sequence acts as a microtubule depolymerase, its activity is reduced relative to MCAK. By contrast the ancestral Kinesin-13 motor domain is the fastest microtubule depolymerase studied to date, with a depolymerisation rate more than an order of magnitude faster than MCAK.

## Results

### An ancestral Kinesin-13 is the fastest microtubule depolymerase observed to date

To comprehensively sample the sequence space occupied by the motor domain of a kinesin family, we generated reference sequences for the motor domain of the microtubule depolymerising Kinesin-13 family from an alignment of the motor domains present in a diverse set of eukaryotes^10^. This motor domain set encompasses the widely-distributed 13A subfamily (including human KIF24 and Kinesin-13 sequences from plants, fungi and diverse protozoa) and the 13B subfamily, which is restricted to animals and close relatives (and includes the well-studied MCAK/KIF2C in addition to KIF2A and KIF2B). We created two representative motor domain sequences: 1) the ‘consensus’ sequence (Con13), containing the most common residue found at every sequence position in the Kinesin-13 family, and 2) the ‘ancestral’ sequence (Anc13), which is the sequence from which all extant Kinesin-13 proteins are inferred to have descended (*i.e*. the last common ancestor for the family; Fig 1a). We expressed and purified protein constructs containing these motor domain sequences with the addition of the neck region of the human Kinesin-13, MCAK, and EGFP for visualisation (Fig 1b). The MCAK neck is included as it is shown to be required for maximal depolymerisation activity of the MCAK motor domain^28, 29^. These reference sequences share 81% identity with each other and ~75% identity with human MCAK across the motor domain, but do not match the sequence of any identified extant Kinesin-13 (SI Fig 1). The differences between these reference motor domains and MCAK are not clustered, but spread throughout the motor domain, with sequence changes found in all major elements of secondary structure (Fig 1c).

**Figure 1:**
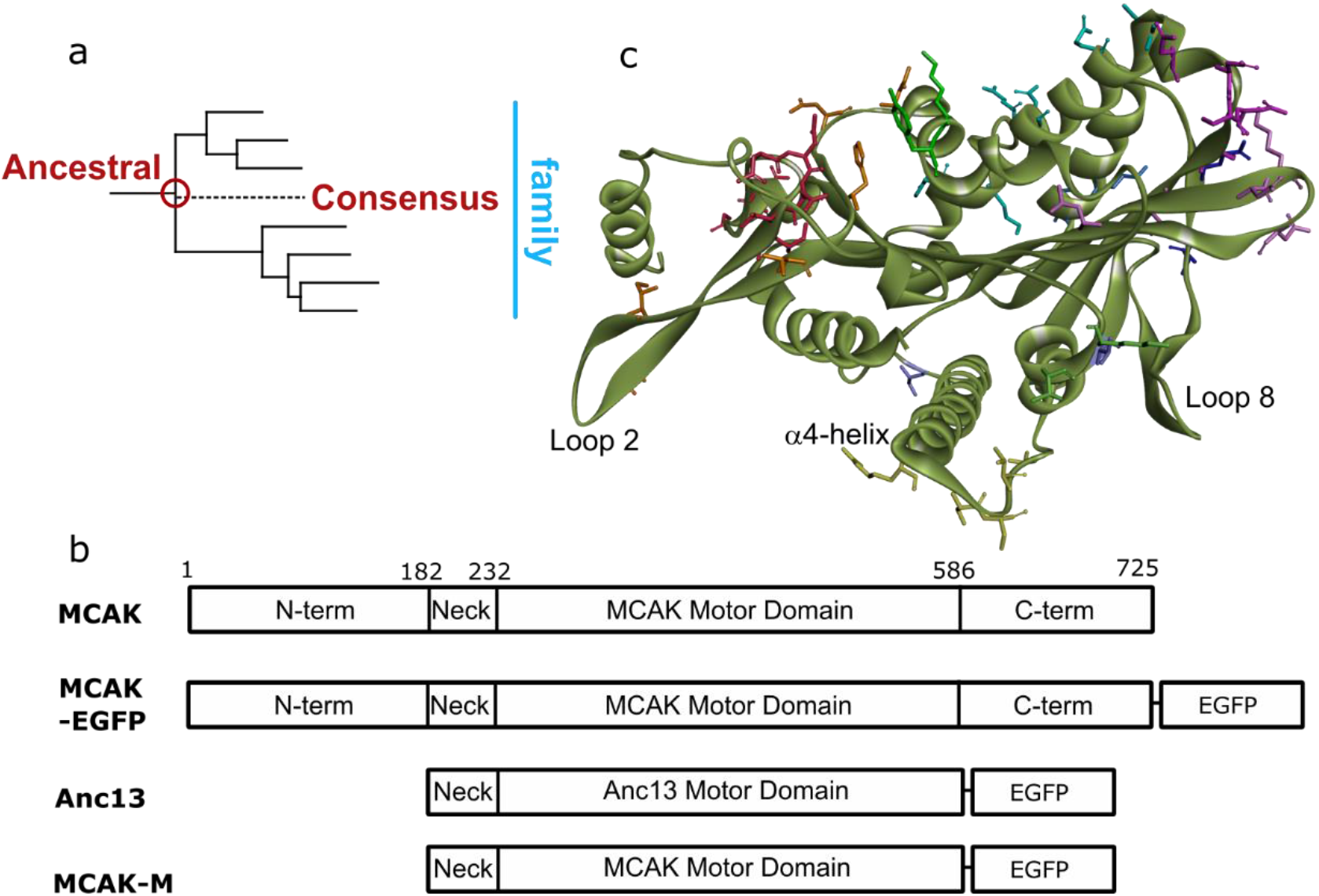
Creation of a Kinesin-13 family ancestral motor domain. a) Location of the ancestral-13 (Anc13) motor domain sequence on the Kinesin-13 family phylogenetic tree. b) Ribbon diagrams of the expression constructs used in this work: full-length MCAK (MCAK), full-length MCAK with C terminal EGFP tag (MCAK-EGFP), the ancestral-13 construct (Anc13) the human MCAK neck region is included at the N-terminal end of the Anc13 motor domain, a version of the Anc13 construct containing the MCAK motor domain in place of the Anc13 motor domain (MCAK-M). c) Prediction of the structure of the Anc13 motor domain with the side-chains of residues that differ from the sequence of human MCAK shown in stick form.

We measured the depolymerisation activity of these Kinesin-13 reference sequences, using GMPCPP stabilised fluorescently-labelled microtubules (Fig 2). Both the consensus and ancestral constructs depolymerised microtubules. The depolymerisation rate of the consensus motor was 0.67±0.28 μm/min, which is ~33% of the rate observed for full-length human MCAK and ~50% of EGFP-labelled full-length MCAK (Table 1). The ancestral motor depolymerised microtubules at 23.05±5.23 μm/min, 11-fold faster than full length MCAK and 17-fold faster than EGFP-labelled MCAK (Fig 2a-c). These data make the inferred ancestral Kinesin-13 motor domain the fastest microtubule depolymerase observed to date.

**Figure 2:**
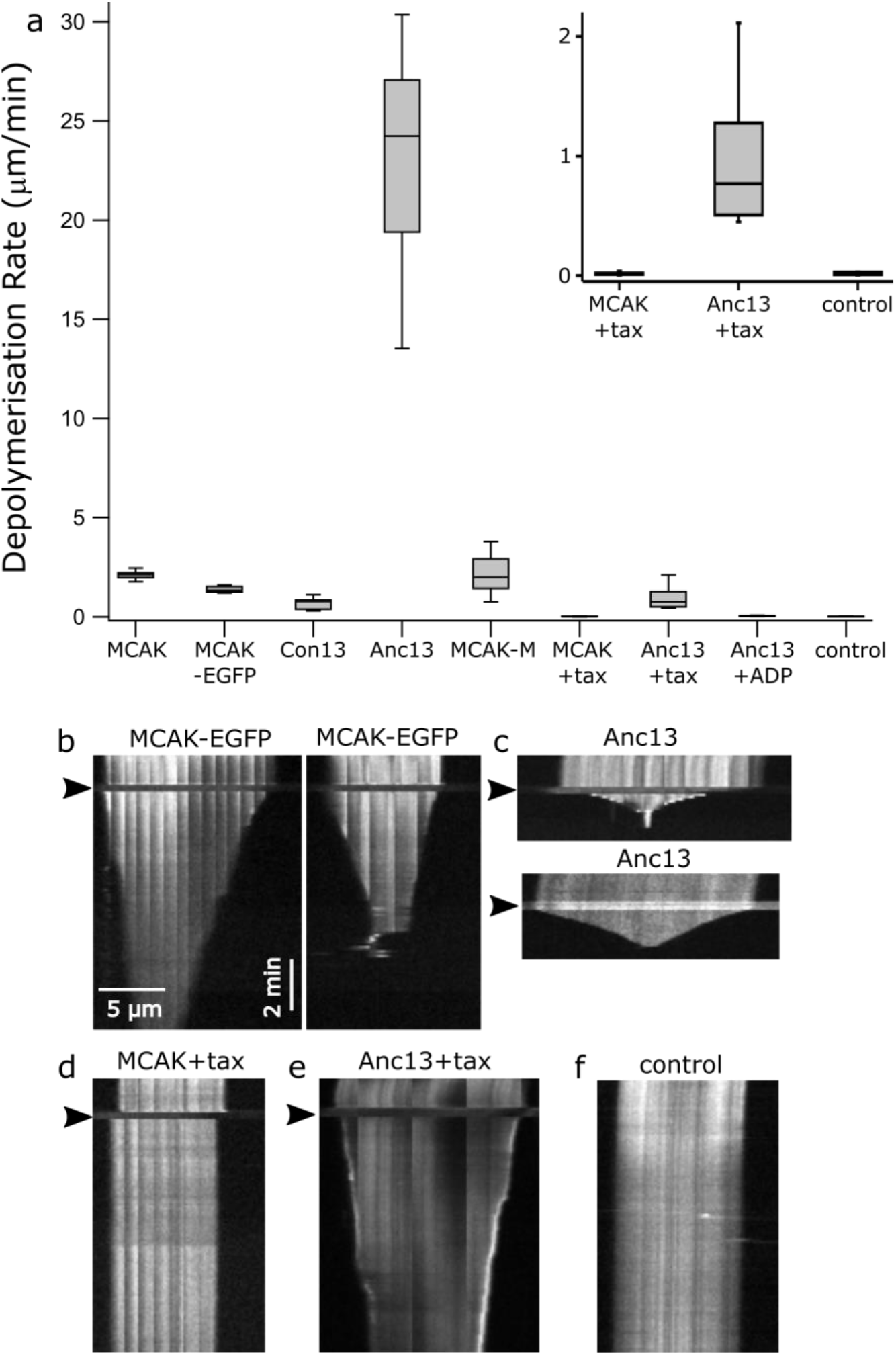
Ancestral-13 has extremely potent microtubule depolymerising activity. a) Microtubule depolymerisation rates for human MCAK and MCAK-GFP compared with Con13, Anc13, MCAK-M, MCAK-GFP, Anc13 in the presence of 1mM taxol, Anc13 in the presence of 1mM ADP rather than ATP and a no added protein control. *Inset:* Zoom in on plots for MCAK and Anc13 in the presence of taxol compared with the basal microtubule depolymerisation rate. b-f) Representative kymographs showing the activity on individual microtubules of b) MCAK-GFP, c) Anc13, d) MCAK-GFP + 1mM taxol, e) Anc13 + 1mM taxol and f) a microtubule with no added protein (control) showing the basal rate of depolymerisation of GMPCPP stabilised microtubules. The arrowheads indicates the time at which the specified protein was added.

**Table 1:**
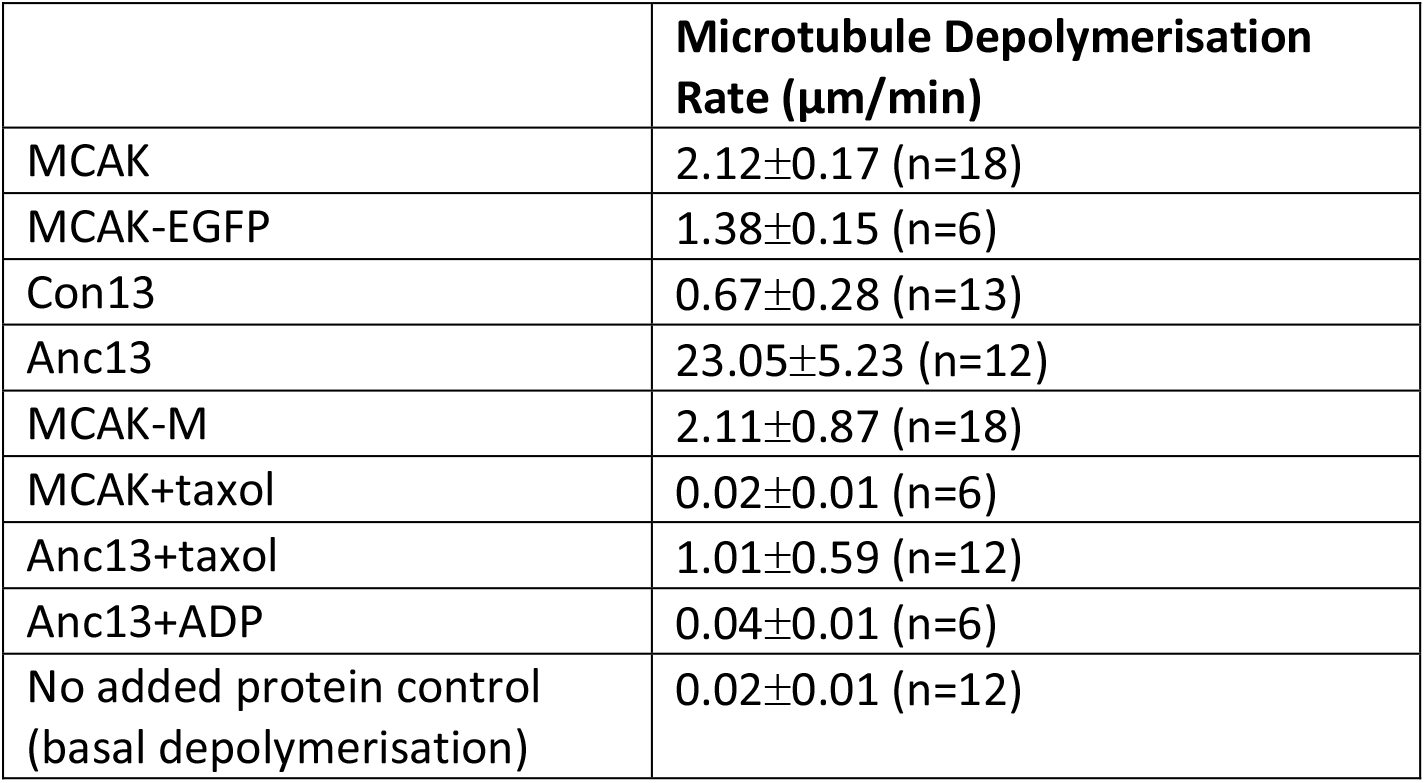
Depolymerisation rates measured for the Kinesin-13 constructs used in this study given as mean ±standard deviation. The basal depolymerisation rate of GMPCPP stabilised microtubules in the absence of any added protein is also shown.

Our investigations of Anc13 show that it requires ATP for microtubule depolymerising activity, as is the case for all members of the Kinesin-13 family studied to date^14, 30^. In the presence of 1mM ADP Anc13 did not depolymerise microtubules (Fig 2a). The depolymerisation rate with ADP as the available nucleotide was 0.04±0.01 μm/min (Table 1), which is similar to the basal depolymerisation rate, ~575-fold lower than Anc13 and over 50-fold lower than MCAK in the presence of ATP.

Full-length MCAK is a homodimer with dimerization occurring through the N- and C-terminal domains which flank the centrally located Kinesin-13 motor domain^31^. The Anc13 construct does not contain the regions of full-length MCAK required for dimerization (Fig 1b) and so is predicted to exist as a monomer. To determine if the increased depolymerisation rate is due to the monomeric nature of the Anc13 motor domain rather than sequence changes within the motor domain, we created the MCAK-M construct in which the ancestral motor domain is replaced with the motor domain of human MCAK in the Anc13 construct (Fig 1b). The microtubule depolymerisation rate for this monomeric version of the MCAK motor domain is 2.11±0.87 μm/min (Fig 2a and Table 1), not significantly different to full-length dimeric MCAK (p=0.96) and ~11-fold lower than Anc13. These data indicate that rather than differences in the oligomerisation state between Anc13 and full-length MCAK, the increased rate of microtubule depolymerisation for Anc13 is due to sequence changes in the motor domain.

### Ancestral-13 depolymerises highly stable microtubules which are impervious to MCAK

Full-length MCAK can depolymerise microtubules stabilised either by incorporation of the slowly hydrolysable GTP analogue GMPCPP or by binding of the drug Taxol, but does not depolymerise microtubules stabilised by both GMPCPP and Taxol (Fig 2a and d)^27^. Double-stabilised microtubules in the presence of MCAK depolymerise at 0.02±0.01 μm/min (Table 1), not significantly different to the basal rate of depolymerisation in the absence of added protein (Fig 2a, d and f). Anc13, however, depolymerised double-stabilised microtubules at 1.01±0.59 μm/min (Fig 2a and e), 50-fold faster than the basal depolymerisation rate and ~48% of the depolymerisation rate of single-stabilised microtubules by full-length MCAK. These data show that the Anc13 motor is a more powerful depolymerase than MCAK, causing significant depolymersation of microtubules stabilised by conditions which MCAK cannot overcome.

### The ancestral Kinesin-13 has increased microtubule affinity, but does not specifically distinguish the microtubule end

To understand how Anc13 exhibits such potent microtubule depolymerising activity, we observed the interaction of Anc13 with microtubules using single molecule TIRF microscopy (Fig 3). The affinity for microtubules of Anc13 is much higher than MCAK, single molecules of MCAK on microtubules are observed at low nanomolar concentrations, whereas for Anc13 we found that picomolar concentrations were required. The data shown in Fig 3a were collected at 8nM MCAK, whereas the data shown in Fig 3b were collected at 40pM Anc13. We calculated a rate of attachment to the microtubule of 62.4±21.5 nM^−1^ s^−1^μm^−1^ for Anc13, which is ~100-fold higher than the microtubule on-rate previously determined for MCAK (Table 2)^12, 16^.

**Figure 3:**
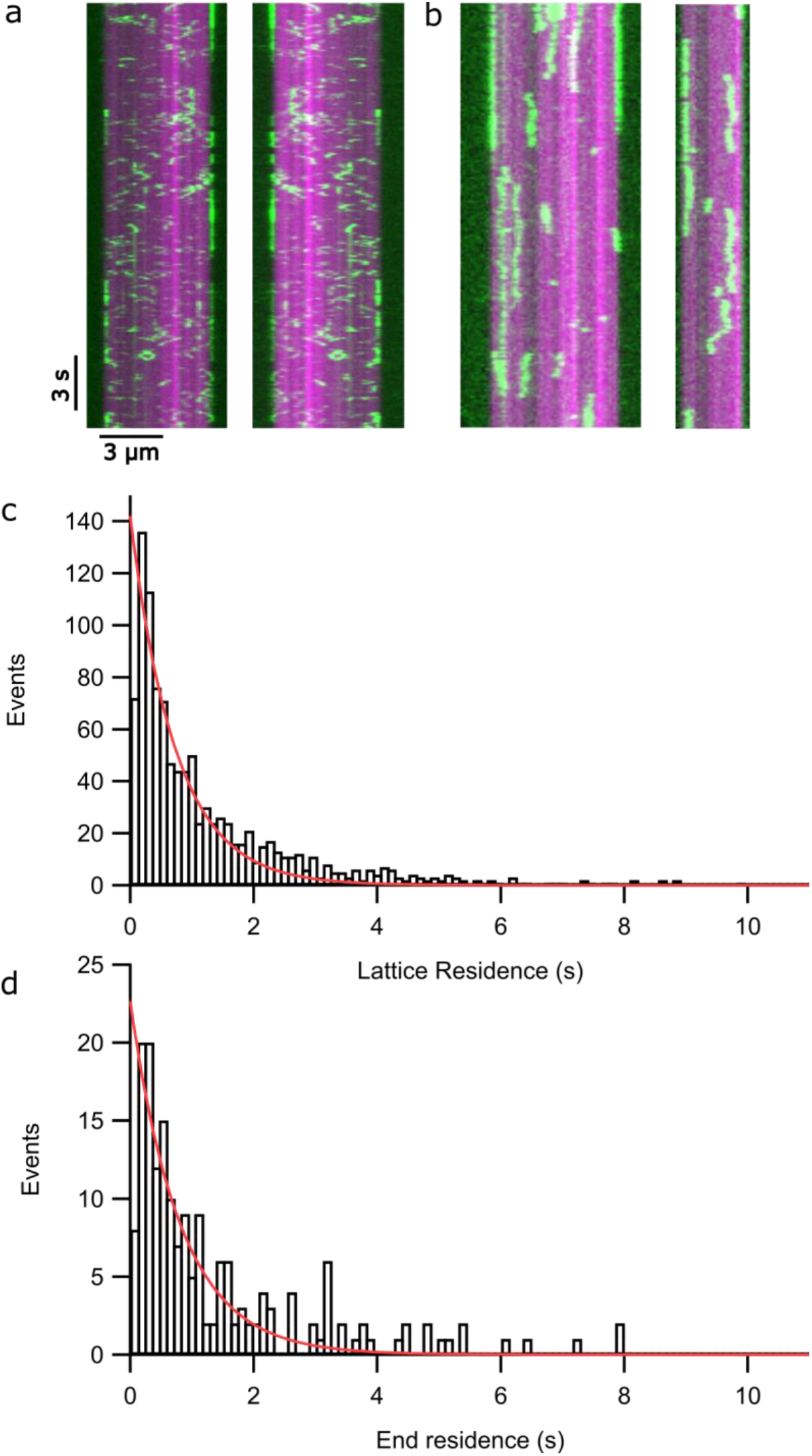
Ancestral 13 does not discriminate the microtubule end from the lattice. a, b) Kymographs showing the interaction of a) MCAK-EGFP b) Anc13 with with GMPCPP-stabilised microtubules (magenta). c, d) Number of events of a particular residence time for Anc13 on the c) microtubule lattice and d) microtubule end.

**Table 2:** Microtubule on and off rates for MCAK and Anc13, given as mean ±standard deviation, calculated using data obtained from single molecule TIRF microscopy assays.

| | $k_{\text{on}}$ ($\mu\text{m}^{-1} \text{s}^{-1} \text{nM}^{-1}$) | $k_{\text{off}}$ MT lattice ($\text{s}^{-1}$) | $k_{\text{off}}$ MT end ($\text{s}^{-1}$) |
| --- | --- | --- | --- |
| Anc13 | $62.4 \pm 21.5$ | $1.36 \pm 0.06$ | $1.23 \pm 0.10$ |
| MCAK | $0.52 \pm 0.34$ | $2.90 \pm 0.16$ | $0.98 \pm 0.06$ |

MCAK displays mostly short (<1s) diffusive interactions with the microtubule lattice. Only at microtubule ends are longer lived interactions observed. Typically, ~30% of MCAK end-binding events last >2s, whereas no lattice-interaction events >2s are observed^16^. In agreement with this, the calculated koff for MCAK when bound to the lattice is ~3-fold higher than at microtubule ends (2.90±0.16 s^−1^ and 0.98±0.06 s^−1^, respectively; Table 2). In contrast, for Anc13 long interaction events are observed on both the microtubule lattice and end, 23% of end-binding events and 20% of lattice binding events are >2s (Fig 3b). The off rates for the microtubule lattice and microtubule end, calculated for Anc13 are similar, k_off_ (lattice) = 1.36±0.06 s^−1^ and k_off (end)_ = 1.23±0.10 s^−1^. The ratio of k_off (lattice/end)_ for Anc13 is 1.11 ±0.14, not significantly different from 1, indicating that Anc13 does not discriminate between the microtubule end and the lattice.

### Ancestral Kinesin-13 has a high basal rate of ATP turnover which is relatively insensitive to the presence of microtubules

To understand how the turnover of ATP is coupled to microtubule depolymerisation by Anc13, we measured ATP turnover in solution, in the presence of unpolymerised tubulin and in the presence of GMPCPP-stabilised microtubules (Table 3). The solution/basal ATPase for Anc13 was 0.44±0.04 s^−1^, which is ~200-fold higher than for full-length MCAK (Table 3). This high rate of ATP turnover in the absence of tubulin could be due to the lack of the N- and C-terminal domains, which have been shown to interact with the motor domain of MCAK and may inhibit ATP turnover^32, 33^. However, the basal ATPase for MCAK-M, the MCAK motor domain lacking the N- and C-terminal regions, was 0.003±0.001s^−1^, not significantly different from full-length MCAK (p=0.25).

**Table 3:**
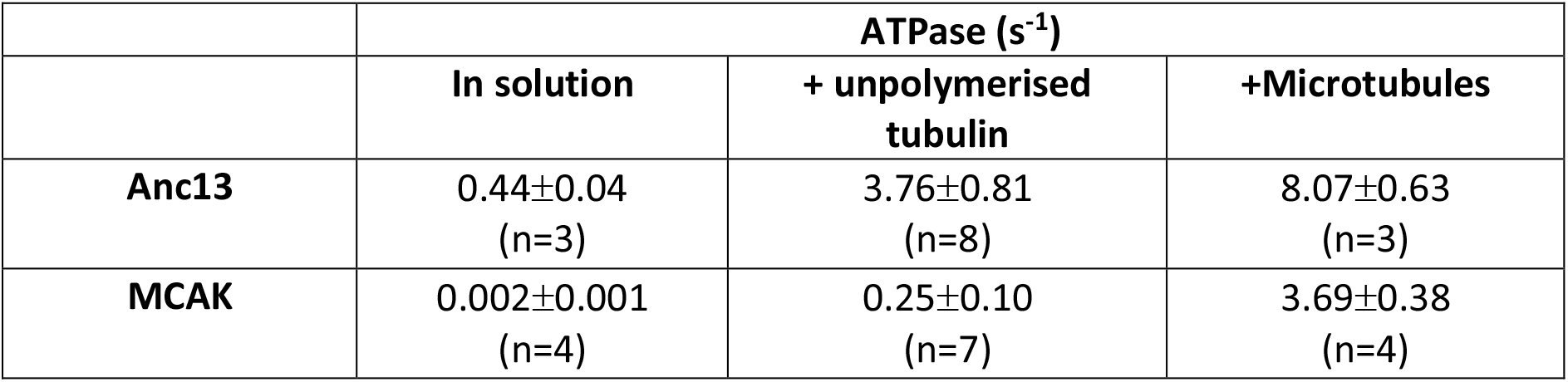
ATPase rates for MCAK and Anc13, given as mean ±standard deviation, measured in solution (basal rate in the absence of tubulin), in the presence of unpolymerised tubulin and in the presence of microtubules.

ATP turnover by Anc13 was accelerated by unpolymerised tubulin. The ATPase rate for Anc13 in the presence of 10μM tubulin was 3.76±0.81s^−1^, an increase of ~8-fold over the basal ATPase. However, this fold increase in ATP turnover is small compared with MCAK for which unpolymerised tubulin causes a ~100-fold increase over the basal rate of ATP turnover^27^.

The fold acceleration in ATP turnover caused by the presence of microtubules is also small for Anc13 compared with MCAK (Table 3). Whilst the microtubule-stimulated ATPase for MCAK is accelerated ~2000-fold with respect to the basal ATPase, the microtubule-stimulated ATPase for Anc13 is 8.07±0.63 s^−1^, only a ~20-fold increase over the basal ATPase.

These data show that the ancestral Kinesin-13 construct has a high rate of ATP turnover even in the absence of tubulin, whereas the MCAK motor domain placed in the same context, as part of the MCAK-M construct, has a very low basal ATPase similar to full-length MCAK. This indicates that the high basal ATPase of Anc13 is an intrinsic property of this motor domain, rather than a result of the absence of the N- and C-terminal regions that flank the motor domain in full-length MCAK.

### The Ancestral-13 motor domain, placed within a full-length Kinesin-13, retains potent depolymerisation activity and promotes internal breaking of microtubules

To understand the behaviour of the ancestral-13 motor domain within the context of a full-length extant Kinesin-13, we created a construct in which the motor domain of full-length wild-type MCAK was replaced with the ancestral-13 motor domain, MCAK-Anc13 (Fig 4a). This construct depolymerised GMPCPP-stabilised microtubules at 19.50±4.87 μm/min, ~9-fold faster than the depolymerisation rate for MCAK and not significantly different (p=0.1) from the depolymerisation rate observed for the Anc13 construct (Fig 4b and c). These data indicate that the ancestral-13 motor retains ultra-rapid depolymerisation activity when placed in the context of full length MCAK, the addition of surrounding N- and C-terminal regions does not significantly alter its microtubule depolymerisation activity.

**Figure 4:**
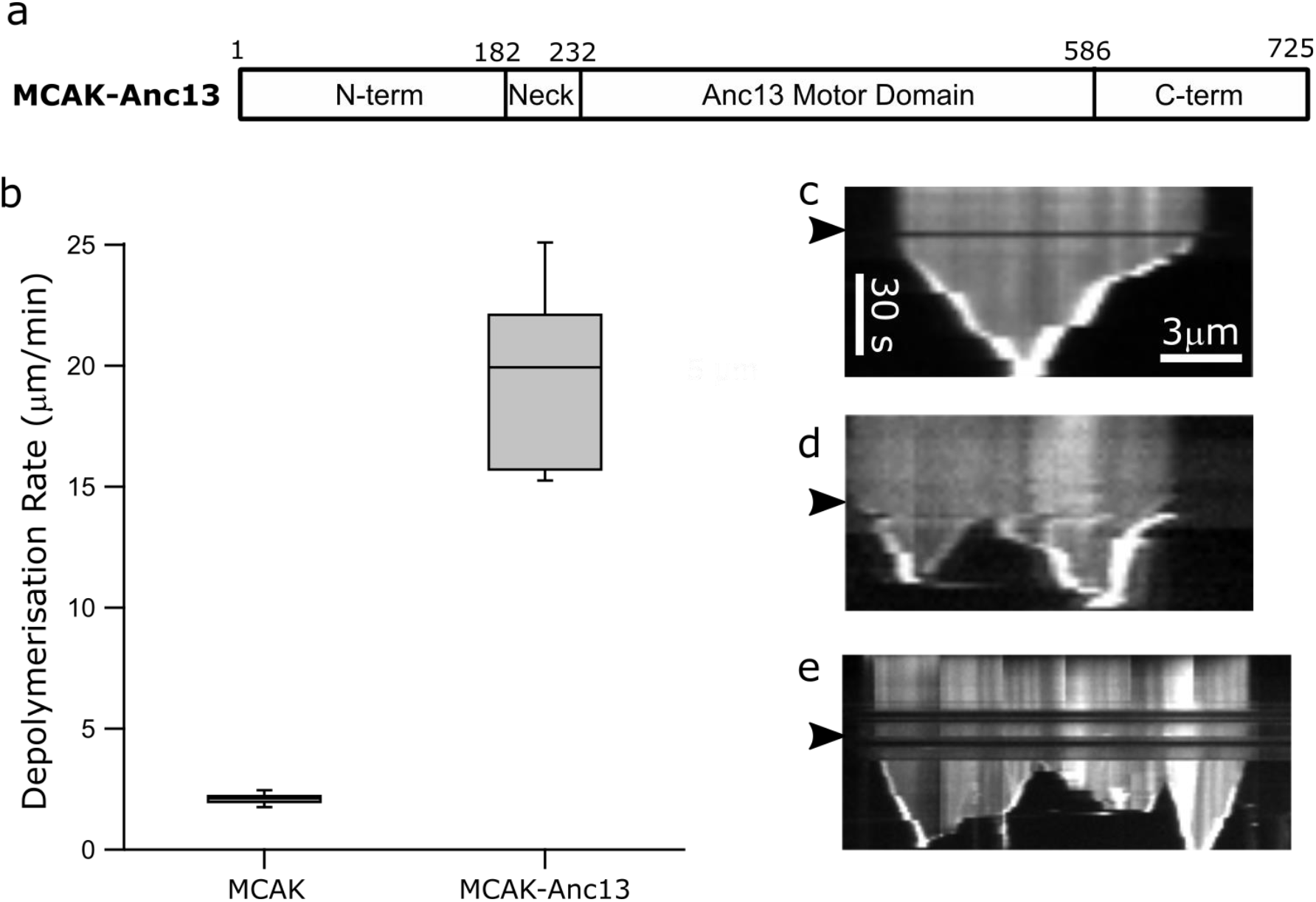
The ancestral-13 motor domain in the context of full-length MCAK has extremely potent microtubule depolymerising activity. a) Ribbon diagram of the expression construct in which the ancestral-13 motor domain is placed within full-length MCAK. Residues 1-232 and 586-725 are the sequence of WT-MCAK. b) Microtubule depolymerisation rates for MCAK and MCAK-Anc13. c-e) Representative kymographs showing the activity on individual microtubules of MCAK-Anc13. d-e) Kymographs showing examples of microtubule which break and continue to depolymerise from the new internally generated ends. The arrowheads indicate the time at which MCAK-Anc13 was added.

In contrast to both MCAK and Anc13, the MCAK-Anc13 construct was observed to promote breaking of microtubules leading to depolymerisation from internal sites (Fig 4d and e). This activity is not frequently observed when microtubules are subjected to depolymerisation by MCAK, which typically causes depolymerisation only from microtubule ends. Quantification of the frequency of internal breaking for microtubules longer than 5μm showed that in the presence of MCAK ~5% of microtubules (n=58) broke at internal sites, whilst in the presence of MCAK-Anc13 ~70% of microtubules (n=47) broke and began to depolymerise from these breaks at a similar rate as from the original microtubule ends. These data indicate that the ancestral motor is an extremely potent depolymerase both as a synthetic monomeric construct and as a dimer in the context of a full-length Kinesin-13.

## Discussion

### The ancestral Kinesin-13 motor domain is a powerful, ultra-rapid microtubule depolymerase

The design and study of reference sequences for the Kinesin-13 motor domain provides insight into the evolution of the Kinesin-13 family and generalised information on the activity of the family as a whole. Both the consensus and ancestral motor domains have microtubule depolymerising activity and depolymerise microtubules from both ends. The consensus sequence (Con13), which contains the most common residue at each position across the family, depolymerises microtubules at ~1/3 the rate of our comparator MCAK. This agrees with the assumption that all members of the Kinesin-13 family are microtubule depolymerases.

In contrast to the consensus protein, the microtubule depolymerisation rate for the ancestral Kinesin-13 motor domain (Anc13) is an order of magnitude faster than the most highly studied member of the Kinesin-13 family, MCAK. To our knowledge, this synthetic protein is the fastest microtubule depolymerase observed to date. In fact, Anc13 has such potent activity that it can depolymerise microtubules under stabilising conditions in which MCAK has no effect. These observations suggest that the founding members of the Kinesin-13 family had much more potent depolymerase activity than Kinesin-13s currently existing in nature (or at least those studied to date). Of course, this activity may have been tempered in ancestral cells by tight regulation of expression or localisation, or by inhibition by regions adjacent to the motor domain that can no longer be reconstructed by phylogenetic analysis. However, potent depolymerase activity is retained when the ancestral-13 motor domain is placed in the context of full-length MCAK surrounded by the non-motor N- and C-terminal regions. It appears that Kinesin-13s in extant lineages have not evolved to become ‘better’ depolymerisers but have instead evolved away from ultra-rapid ‘super’ depolymerase activity toward milder action.

### Ultra-rapid depolymerisation is associated with inability to focus activity to microtubule end

Single molecule TIRF data shows that, whilst Anc13 has higher overall affinity for microtubules, it does not distinguish between the microtubule end and the microtubule lattice. Rather, Anc13 displays long binding events at all positions on the microtubule. This contrasts with MCAK for which long binding events are observed only at or near microtubule ends^16^, focusing activity to the place in which tubulin removal can most easily occur and protecting the microtubule body from destabilisation. The ultimate extension of this lack of ability to focus activity to the microtubule end is observed for the MCAK-Anc13 construct, which causes internal breaks within microtubules and promotes depolymerisation not only from the microtubule end, but also from these break sites. This type of behaviour is rarely observed for extant kinesin-13s but resembles the activity of known microtubule severing proteins such as katanin^34^.

### Ancestral Kinesin-13 removes multiple tubulin dimers per ATP consumed

The stoichiometry of ATP consumed per tubulin removed by MCAK has previously been measured to range from 1 ATP to >20 ATPs per tubulin^27^. During processive depolymerisation of GMPCPP stabilised microtubules MCAK removes on average 1 tubulin for each ATP consumed: every ATPase cycle is productive for depolymerisation. With progressively more stable microtubules MCAK fails to remove tubulin for some ATP turnover cycles. Thus the proportion of productive cycles decreases from 100% for GMPCPP stabilised microtubules, to 50% for taxol stabilised microtubules and to <5% for double stabilised microtubules^27, 30^. However, from currently available data, there is no indication that MCAK removes more than 1 tubulin dimer on average per ATP under any conditions.

The observation that the microtubule-stimulated ATPase for Anc13 was only ~2-fold higher than MCAK whilst the microtubule depolymerisation rate was >10-fold higher was unexpected, as this indicates that Anc13 removes more than a single tubulin for each cycle of ATP turnover. The stoichiometry of ATP consumed per tubulin removed for Anc13 is ~0.2 ATP per tubulin, indicating that for each ATP consumed Anc13 removes multiple tubulin dimers. From our data we calculate that Anc13 removes on average 5.8±1.4 tubulins per ATP, when acting on GMPCPP stabilised microtubules.

To explain these data, Anc13 must be able to bind at locations prior to the terminal tubulin dimer and destabilise tubulin subunits relatively distant to the microtubule end (Fig 5). Our data indicate that on average 6 tubulin dimers are removed per single cycle of ATP turnover. This could mean that Anc13 on average binds 6 dimers away from the microtubule end and causes detachment of the bound tubulin and the remaining subunits from the same protofilament (Fig 5b) or, perhaps more likely, that Anc13 binds anywhere up to 6-dimers away from the microtubule end and causes destabilisation of the bound tubulin leading to detachment of the remaining subunits from the same protofilament plus tubulin subunits from neighbouring protofilaments, causing on average 6 dimers to detach from the microtubule (Fig 5c).

**Figure 5:**
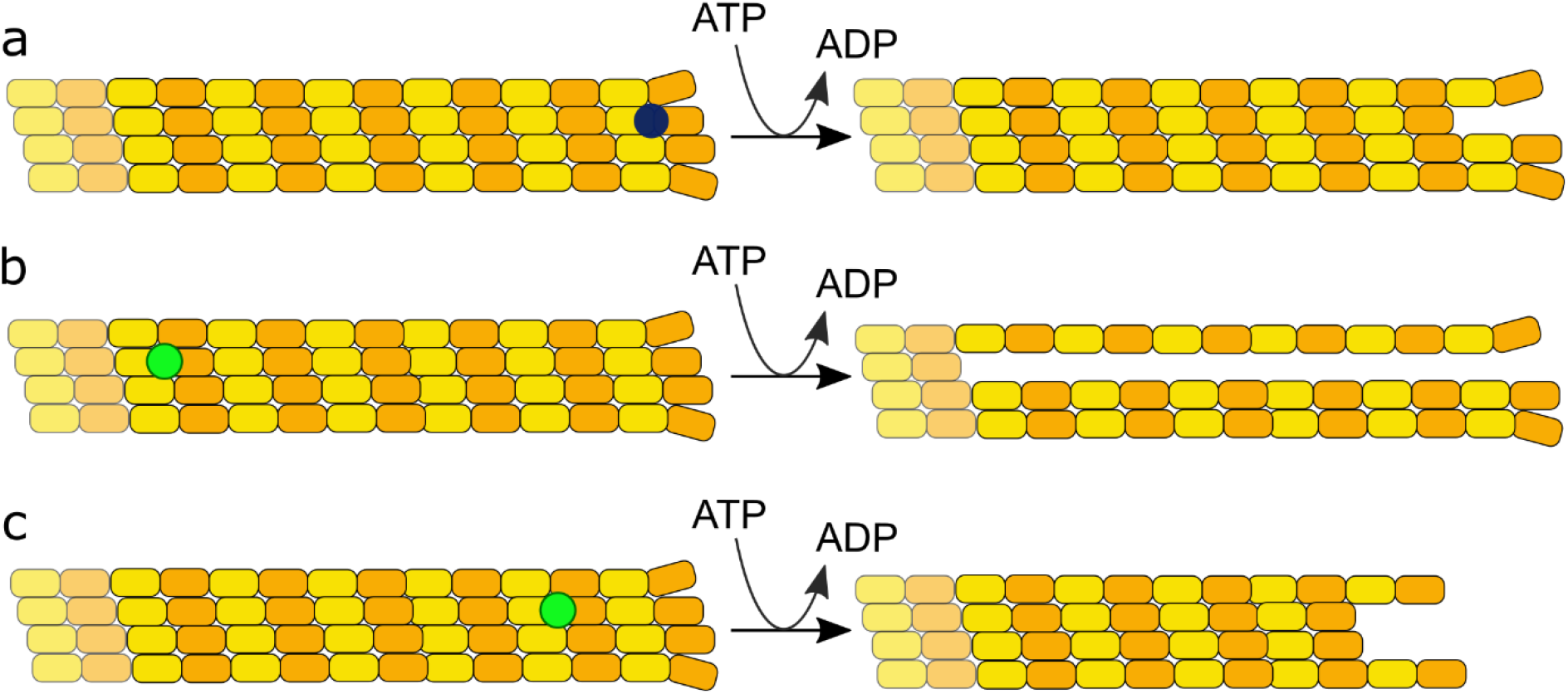
Microtubule depolymerisation by MCAK compared with possible mechanisms of depolymerisation by Anc13. Schematic of a microtubule depolymerised by a) MCAK and b & c) Anc13. a) Available data indicates that MCAK typically binds to and removes only the terminal tubulin dimer. The highest ratio of tubulin removed to ATP consumed measured for MCAK is 1:1. MCAK on average removes a single tubulin for each ATP consumed. b & c) Our data shows that Anc13 removes on average 5.8±1.4 tubulin dimers per ATP consumed. This suggests that Anc13 binds to and destabilises tubulin dimers prior to the terminal protofilament dimer. b) Anc13 could bind to the 6th dimer prior to a protofilament end and cause detachment of the rest of the same protofilament or c) Anc13 could bind closer to the protofilament end and cause detachment of tubulin subunits from the same protofilament plus tubulin from neighbouring protofilaments.

Tubulin dimers farther from the microtubule end are likely more stably embedded within the microtubule than those close to the end. Therefore, to destabilise tubulin subunits at sites prior to the terminal dimer of a microtubule protofilament would require higher affinity binding to deform tubulin dimers located at more stable positions within the microtubule. Anc13 not only binds microtubules with higher affinity than MCAK, but also can depolymerise microtubules under stabilising conditions that cannot be overcome by MCAK. Further, when the ancestral-13 motor domain acts as a dimer in the context of full length MCAK, it causes breaking of microtubules at internal positions. These data indicate that the ancestral-13 motor domain can promote deformation of tubulin subunits distant to the microtubule end resulting in removal of multiple dimers per ATP turnover cycle and internal breaking of microtubules, resulting in the most rapid microtubule depolymerase activity observed to date.

### The Kinesin-13 family evolved away from potent activity toward more controlled microtubule depolymerisation

Our data suggest that the founding members of the Kinesin-13 family were highly potent, ‘super’ depolymerases. Comparison of the activity of a phylogenetically inferred ancestral Kinesin-13 motor domain with extant Kinesin-13s implies that as the family developed its members evolved away from rampant, depolymerisation toward more controlled, but less rapid activity. Comparison of the biochemical parameters for Anc13 with those previously determined for MCAK, shows that whilst Anc13 can depolymerise microtubules an order of magnitude faster, this comes at the cost of efficiency in terms of ATP turnover and specificity in localization.

It is not obvious which sequence changes between Anc13 and MCAK are responsible for the increased potency of the ancestral-13 motor domain. Sequence changes are scattered throughout the motor domain (SI Fig S1) and no sequence alterations are found in any of the conserved kinesin specific nucleotide binding motifs or in the Kinesin-13 family specific microtubule binding motifs.

In comparison to the ancestral sequence, modern-day MCAK has gained several characteristics that are presumably beneficial for the physiological roles of extant members of the Kinesin-13 family. Mitotic spindle and flagellar length control are proposed to be the principal roles of Kinesin-13s conserved throughout the evolution of eukaryotes^35^. Highly potent, ultra-rapid depolymerisation that is not focused at microtubule ends, but is ‘katanin-like’ causing internal breaking of microtubules, is likely not beneficial for these roles which require precise regulation of microtubule length. Our data suggests that better regulated activity has outcompeted rapid but less focused depolymerisation activity during the evolution of the Kineisn-13 family. The design and creation of bioinformatically-inferred reference sequences provides insight into the evolution of the Kinesin-13 family and could provide similar insight into other kinesin families or other protein families. The approach described allows comprehensive sampling of sequence space, which could provide insight into the evolution of function for any large paralogous protein family.

## Methods

### Inference of reference motor domain sequences

Reference sequences for the Kinesin-13 family were based on the previous analysis of 1624 kinesins proteins encoded in the genomes of 45 diverse extant eukaryotes^10^. The alignment of the motor domains and the phylogenetic tree for the Kinesin-13 family were extracted from this larger analysis of the full kinesin superfamily. The Kinesin-13 subtree encompasses 92 protein sequences from 38 eukaryotes (i.e. all those in the analysis that encode a Kinesin-13 family member). To avoid ancestral reconstruction being heavily biased by divergent sequences, the 13C subfamily (which includes divergent Kinesin-13-like sequences from various microbial eukaryotes) was excluded and only subfamilies 13A and 13B considered (75 sequences in total).

Consensus motor domain sequence was inferred by simple most-common residue method, with ties resolved based on most likely residue by BLOSUM62 score. Ancestral sequences were inferred by maximum likelihood using the WAG substitution matrix as implemented by fastML v3.1 (--seqType aa --SubMatrix WAG --indelReconstruction ML^36^). A single internal polytomy in the 13B subfamily tree was resolved based on branching of the included species, although this has no impact on inference of the ancestral state of the 13A/B family.

### Expression and purification of kinesin reference constructs

Reference protein sequences were reverse translated using a *Spodoptera frugiperda* (Sf9) codon usage table (IDT <u>Codon Optimization Tool</u>, www.idtdna.com). These coding sequences were incorporated into derivatives of pFastBac1 encoding the neck region from Hs MCAK on the N-terminal side and eGFP and TEV-cleavable streptavidin-binding peptide- and His-tags on the C-terminal side (see Supplemental Files 1 and 2 or full sequences). Formation of the expected chimeric sequence was confirmed by Sanger sequencing covering the full CDS.

The proteins were expressed in Sf9 cells (Bac-To-Bac expression system; Invitrogen) and purified using Ni-affinity followed by Strep-Tactin affinity purification. Cells were lysed by incubation at 4°C for 1h in 50mM Tris pH7.5, 300mM NaCl, 10% glycerol, 5mM MgCl_2_, 0.1% Tween 20, 10mM imidazole, 1mM DTT and protease inhibitors. The proteins were purified by Ni-affinity chromatography (Ni-NTA agarose, Qiagen) followed by StrepTactin affinity chromatography (StrepTactin Sepharose high performance, GE Healthcare).

### Microtubules and depolymerisation assays

Porcine brain tubulin was purified from homogenised brain tissue using the high ionic strength method^37^. Microtubule depolymerisation rates were determined by measuring the length of immobilised, GMPCPP-stabilised, 25% rhodamine labelled microtubules over time after the addition of 40 nM Kinesin-13 as described previously^16^. The depolymerisation data for MCAK-Anc13 was captured by imaging every 2s, rather than every 5s as was the case for all other proteins studied.

### Single molecule TIRF microscopy

Rhodamine-labelled, GMPCPP-stabilised microtubules were stuck onto the surface of flow chambers prepared as described previously^38^. MCAK-eGFP or Anc13 at single-molecule concentrations (4-8nM and 40pM, respectively) were added to a microtubule containing flow cell in BRB20 pH 6.9, 75 mM KCl, 1 mM ATP, 0.05% Tween 20, 0.1 mg/ml BSA, 1% 2-mercaptoethanol, 40 mM D-glucose, 40 mg/ml glucose oxidase, 16 mg/ml catalase. Images were recorded using a Zeiss PS1 Super Resolution Microscope. Microtubules were imaged using laser excitation (561nm) and a 579-620nm emission filter. Movies of eGFP-labelled single molecules were collected using TIRF excitation at 488nm and a 495-550 emission filter, 200 imaged were collected per movie with a frame rate of 8.7 Hz.

For analysis, each frame of the GFP dataset was colour-combined with the corresponding image of rhodamine-labelled microtubules in FIJI to enable identification of on-microtubule events. Kymographs for individual microtubules were used to measure the duration of individual GFP localization events at the microtubule end and on the lattice, also in FIJI^39^.

### ATPase assays

ATPase rates in solution were measured using 3 μM kinesin by monitoring the production of ADP using HPLC as described previously^40^. For assays with added tubulin or microtubules, 0.1 μM kinesin was used and the production of ADP monitored by linking it to the oxidation of NADH^16, 41^. For both assays the change in concentration of ADP per second was divided by the concentration of kinesin motor domain to give the ATPase activity per second per active site.

## Supplementary Information

**SI Figure S1:**
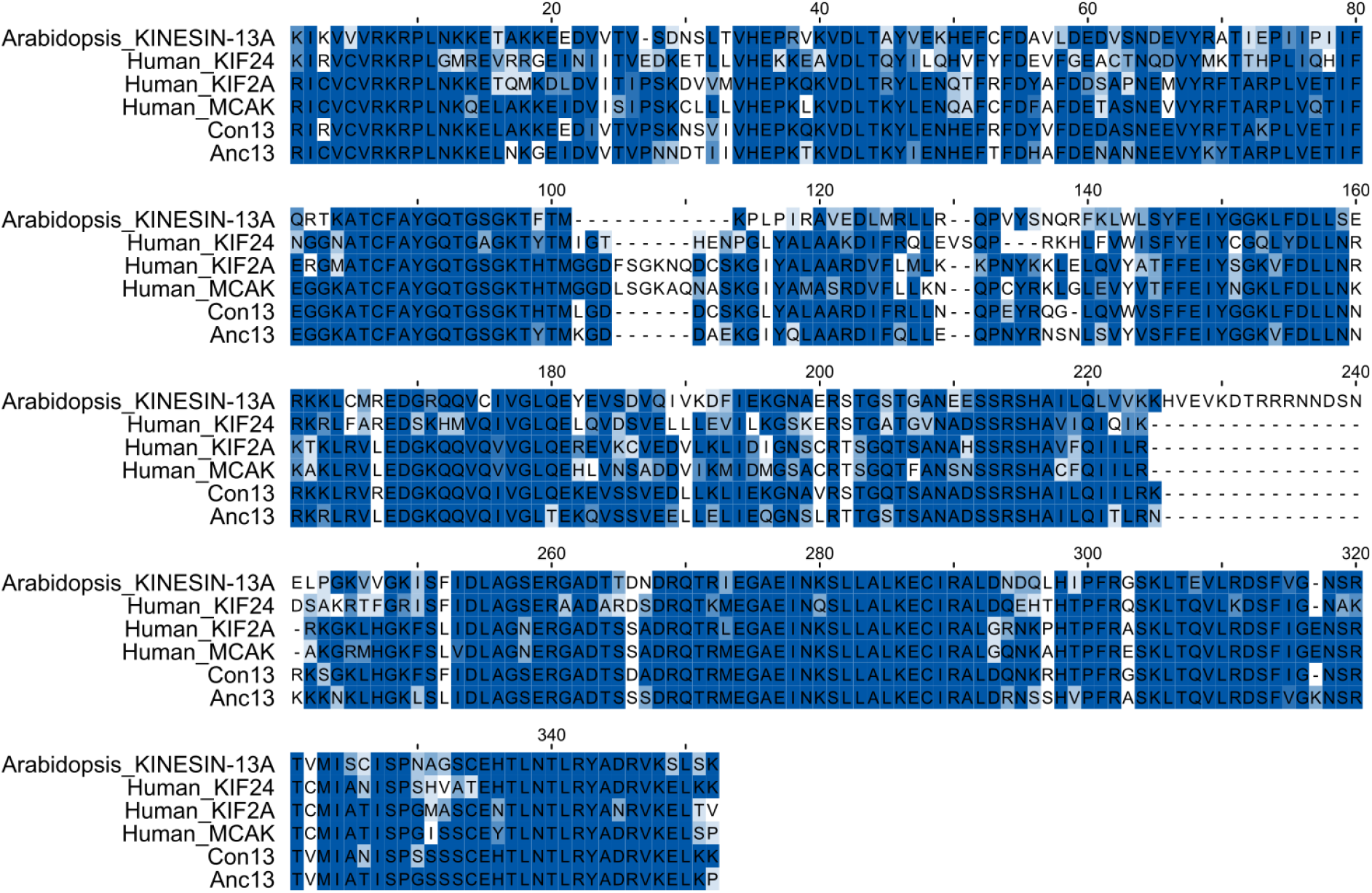
Comparison of the motor domain of members of the Kinesin-13 family. Alignment of motor domain sequences from select members of the Kinesin-13 family is shown together with the Consensus (Con13) and Ancestral (Anc13) Kinesin-13 sequences.

## Notes

### Competing Interest Statement

The authors have declared no competing interest.

